# Prairie management practices influence biodiversity, productivity and surface-atmosphere feedbacks

**DOI:** 10.1101/2025.01.02.631106

**Authors:** Ran Wang, John A. Gamon, Katharine F. E. Hogan, P. Roxanne Kellar, David A. Wedin

**Affiliations:** School of Natural Resources, University of Nebraska-Lincoln, Lincoln, Nebraska, USA; Department of Biological Sciences, Northern Illinois University, DeKalb, Illinois, USA; Department of Biology, University of Nebraska at Omaha, Omaha, Nebraska, USA

**Keywords:** Grassland, restoration, imaging spectroscopy, thermal remote sensing, prescribed fire, controlled burning

## Abstract

Restoration efforts aim to reestablish grassland cover and maintain ecosystem services. However, there is a lack of systematic evaluation of the effects of grassland restoration options on integrated biodiversity, carbon, ecosystem function, and climate feedbacks.

Through a multi-year grassland restoration experiment in a tallgrass prairie site in Nebraska, USA, we investigated how different management practices affected biodiversity, productivity and the surface-atmosphere feedbacks using a combination of in situ measurements and airborne hyperspectral and thermal remote sensing.

Our findings suggested that certain management treatments altered species diversity, productivity and energy balance. Higher diversity plots had higher vegetation cover, albedo, canopy water content and lower surface temperature, indicating clear effects of management treatments on grassland ecosystem processes influencing surface-atmosphere feedbacks of mass and energy.

The coherent responses of multiple airborne remote sensing indices illustrate clear co-benefits of grassland restoration practices that enhance ecosystem productivity and biodiversity and mitigate climate change through surface atmosphere feedbacks, offering a new strategy to address the global challenges of biodiversity loss and climate change in prairie ecosystems.

## Introduction

Grasslands constitute about 40% of the Earth’s terrestrial area and cover a range of geographic locations (White et al., 2000). Besides supporting agricultural production through grazing, healthy grasslands sustain a variety of plant and animal species, stabilize soil and prevent erosion, improve water retention and groundwater recharge, and provide habitat and food sources for pollinators (Bengtsson et al., 2019, Murphy et al., 2016). Grasslands can also be important soil carbon sinks and store approximately one third of global terrestrial carbon (Bai and Cotrufo, 2022). However, grasslands have experienced a worldwide decline during the last century, leaving half of grasslands degraded (Bardgett et al., 2021) with few intact grasslands remaining (Scholtz and Twidwell, 2022). Grassland degradation is often caused by landcover conversion to crops and forestry, land abandonment, overgrazing, woody encroachment and species invasion (Parr et al., 2014, Bardgett et al., 2021). Abandonment of historical indigenous management practices (e.g., frequent fire) has also altered grassland composition and function (Pyne, 2019).

Altered disturbance regimes (e.g., grazing intensity and timing and frequency of fire) and species reintroduction (e.g., seeding) can gradually restore grassland aboveground diversity and function and below ground structure (Buisson et al., 2022), suggesting that effective restoration strategies may have multiple co-benefits.

Nature-based solutions aim to adapt to and mitigate climate change effects while improving sustainable livelihoods, protecting natural ecosystems and biodiversity, and preventing the degradation and loss of natural ecosystems (IUCN, 2016). Restoring grassland coverage and biodiversity can serve as an effective strategy for mitigating negative impacts of global climate change by facilitating relatively fast and resilient carbon sequestration (Fargione et al., 2018, Griscom et al., 2017, Bai and Cotrufo, 2022). Conservation programs, such as the USDA Conservation Reserve Program (CRP) (Allen and Vandever, 2003, Hellerstein, 2017) and British Coronation Meadows (Thomas, 2021), aim to re-establish valuable land cover to help improve water quality, prevent soil erosion, enhance carbon sequestration, reduce nitrogen runoff and provide healthy habitat for wildlife and pollinators. Despite reports of successfully achieving these goals individually (Gelfand et al., 2011, Becker et al., 2022, Gleason et al., 2011), there is a lack of systematic evaluation of the combined effects on integrated biodiversity, carbon, ecosystem function, and climate feedbacks of grassland restoration practices, especially when in conjunction with heterogenous management processes such as prescribed burning, mowing, and grazing. This leaves a gap in our understanding about how management and restoration practices contribute to critical ecosystem processes that may offer climate benefits.

Terrestrial surface energy balance refers to the equilibrium between incoming solar radiation, energy stored in plant biomass, and outgoing energy in the form of energy reflected (albedo) and energy used for heat and evapotranspiration, typically measured as surface temperature (Still et al., 2019, Schulze et al., 2019). Along with mass balance, understanding this energy balance is essential for managing ecosystems, predicting their responses to climate change, and assessing their role in carbon and water cycles (Bonan, 2008). Emerging remote sensing methods can capture states and processes related to mass and energy exchange (Hall et al., 1992), but applying these methods to prairie restoration evaluations remain largely underexplored (Blackburn et al., 2021). One reason is the scale mismatches between common spaceborne remote sensing methods and that of land ownership and management units such as individual fields or pastures that can be readily sampled. Airborne remote sensing, with the combined power of explicit spectral and thermal sampling at fine spatial resolutions, readily matches the scale of typical grassland management practices and offers a powerful set of tools for evaluating the effects of grassland restoration among a range of management treatments.

In this study, we conducted a multi-year experiment in a tall-grass prairie restoration site near Lincoln, Nebraska, USA with various management treatments that included contrasting seeding diversity, burning and mowing practices. Our primary goal was to investigate the impact of several common management strategies on grassland biodiversity (species and functional composition) and ecosystem function (productivity and energy balance). A key objective was to evaluate potential co-benefits of prairie seeding and management treatments with implications for productivity, biodiversity and climate mitigation.

## Materials and Methods

### Field site description and experimental design

This study was conducted at the Bobcat Restoration Experiment (40.71°N, 96.82°W), a ca. 12-ha field located in Lancaster County, Nebraska that has been annually hayed for several decades, partially under United Stated Department of Agriculture (USDA) Conservation Reserve Program (CRP) contract management. Prior to the restoration experiment, the site was dominated by invasive smooth brome (*Bromus inermis* Leyss.) and native Indiangrass (*Sorghastrum nutans* (L.) Nash). The average growing season temperature (May to September) is 28 °C, and the average annual precipitation is approx. 74 cm.

We arranged the treatments into eight blocks of three plots each (24 plots in total; Fig. 1a and Table 1) to evenly distribute seeding treatments and management strategies (haying vs. burning) while accounting for site heterogeneity in soils and extant vegetation. The size of each plot was approx. 0.40 ha (Fig. 1a). The seeding treatments included: (1) a high diversity local ecotype (“high diversity”) seed mix of 154 species from Prairie Plains Resource Institute in Aurora, Nebraska, (2) a local ecotype modified CRP-style pollinator (“mid diversity”) mix of 40 species from Prairie Legacy Inc, Nebraska, and (3) a low-diversity control group with no seed applications. For the high and mid diversity plots, we applied two glyphosate treatments using a boom sprayer (at the standard industry rate of 0.75 lbs/ac) to weaken the pre-existing vegetation, mostly smooth brome (*B. inermis*), on November 18, 2018 and April 19, 2019. The high diversity mix was broadcast from an ATV spreader on April 29, 2019, onto a lightly disked surface to ensure seed-soil contact. The mid diversity mix was drill seeded on May 1, 2019. The control plots received no glyphosate treatments, nor seed applications. Management treatments (Fig. 1a) consisted of annual mowing or haying in August starting 2019 or burning on May 10, 2022. The plots assigned to the haying treatment were mowed during the first growing season of the experiment (July 30^th^, 2019), as not enough biomass was present for effective baling. Haying commenced the following growing season. The detailed seeding and management treatments are further summarized in Tables 1 and 2.

**Fig. 1.**
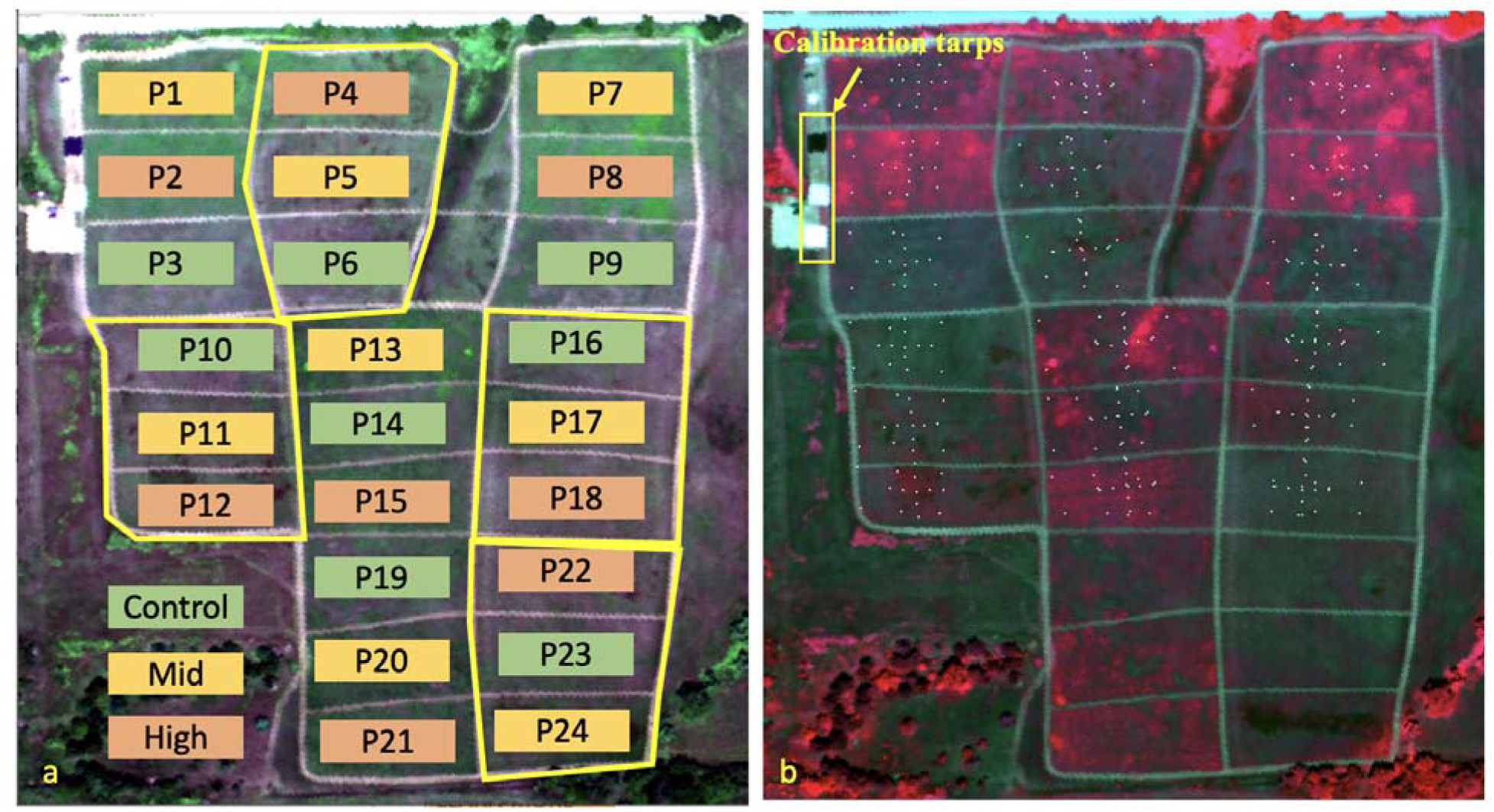
Experimental design of the Bobcat Restoration Experiment. a: Schematic of the plot locations, seeding treatments (control, mid or high diversity) and management (burning and haying) laid on top of the airborne true color image (Red: 680 nm, Green: 550 nm, and Blue: 460 nm). Green plots are control, yellow plots are mid diversity, and orange plots are high diversity. The yellow boundaries indicate blocks that received annual haying since 2019. The rest of the plots were burned on May 10, 2022. b: Locations of the 2022 field vegetation sampling (white dots) on top of the airborne false color image (Red: 800 nm, Green: 680 nm, and Blue: 550 nm). Also shown are the locations of calibration tarps (yellow rectangle). Airborne image was collected using the Kestrel imaging spectrometer on July 20, 2022.

**Table 1.** A summary of management treatments applied to the experimental blocks.

| Experimental block | Plots within block | Management treatment of block | Management date(s) |
| --- | --- | --- | --- |
| 1 | 1,2,3 | Prescribed burn | May 10th 2022 |
| 2 | 4,5,6 | Hay/mow | July 30th 2019; August 10th, 2020; August 20th 2021 |
| 3 | 7,8,9 | Prescribed burn | May 10th 2022 |
| 4 | 10,11,12 | Hay/mow | July 30th 2019; August 10th, 2020; August 20th 2021 |
| 5 | 13,14,15 | Prescribed burn | May 10th 2022 |
| 6 | 16,17,18 | Hay/mow | July 30th 2019; August 10th, 2020; August 20th 2021 |
| 7 | 19,20,21 | Prescribed burn | May 10th 2022 |
| 8 | 22,23,24 | Hay/mow | July 30th 2019; August 10th, 2020; August 20th 2021 |

**Table 2.** A summary of seeding treatments applied to the plots.

| Diversity level | Glyphosate treatments | Seed mix | Seeding application |
| --- | --- | --- | --- |
| Low | NA | NA | NA |
| Mid | Nov 18, 2018; Apr 19, 2019 | Local ecotype modified CRP-style pollinator of 40 species | Drilled |
| High | Nov 18, 2018; Apr 19, 2019 | Local ecotype high diversity seed mix of 154 species | Disc & broadcast |

### Field species data

Species data were recorded for 18 of the 24 plots (Fig. 1b) in August 2022. Within each plot, we started roughly from the center of the plot and laid out four 15-meter long transects in the west-east and north-south directions. We sampled species data at the center of the plot, and at meter 7 and meter 14 along each transect in the four directions (nine locations per treatment plot). In addition to these transects, we also sampled species data from six subplots (in a 2×3 array along the east-west direction; Fig. 1b) within each plot. This design resulted in 15 vegetation sampling positions within each plot. At each sampling location, species percentage cover within a 1m^2^ quadrat was recorded. The six subplots were originally designed to be 5 meters away from the edge of the plot and roughly 30 meters apart from each other. However, the 5-meter margin of the subplots varied over the course of the experiment due to mowing and vehicle traffic around the edges of the plots. To avoid potential edge effects, we removed the data that was sampled within 5 meters from edges of each plot. Using the percentage cover data, we calculated species richness, Shannon’s index based on species, and Shannon’s index based on functional groups (C_3_ grasses, C_4_ grasses, and forbs).

### In-situ proximal remote sensing data

To calibrate airborne data and interpret the treatment effects across the full solar spectral range, we collected ground reflectance data for six plots (P1, P2, P3, P10, P11, P12; Fig. 1) using a portable spectrometer (PSR+ 3500; Spectral Evolution Inc., Lawrence, MA, USA), which covered the 350 to 2500 nm spectral range. Measurements were taken at every meter along the four 15-meter north-south and east-west transects, yielding 60 samples for each plot. Spectral radiance was sampled using a 2-m fiber (Spectral Evolution Inc., Lawrence, MA, USA) at a height of approx. 1.5 m above the ground. The field of view (FOV) of the fiber was 25° and the size of the ground sampling was ∼1 m. A spectralon calibration panel (Labsphere, North Sutton, New Hampshire USA) was measured before field spectral data collection in each plot. Then canopy reflectance was calculated as the ratio between radiances of vegetation and the calibration panel. All the measurements were collected between 10 a.m. and 4 p.m. to minimize shadow effects.

### Airborne data collection and processing

Airborne hyperspectral and thermal images were collected on July 20 2022 (around 13:35 local time) using an imaging spectrometer (AISA Kestrel; Specim, Oulu, Finland) and a thermal imager (IR-TCM 640; Jenoptik Optical Systems GmbH, Jena, Germany) mounted on a fixed-wing airplane (Piper Saratoga; Vero Beach, Florida, USA). Images were collected from approximately 1.5 km above ground level. Air temperature was 30.6 °C at the flight time. Air temperature was measured at the Spring Creek Prairie Site (3.75 km from the field site) within the U.S. Climate Reference Network.

### Hyperspectral data

The Kestrel imaging spectrometer collected visible-near infrared (400 to 1000 nm) hyperspectral data with a 2.3 nm spectral resolution. The FOV is 40° and the ground pixel size was approximately 1m. An inflight GPS and inertial measurement unit (IMU) (RT3000; Oxford Technical Solutions Limited, England) was used to record the location and rotation attributes of the aircraft, which was then used for image geometric correction. To obtain surface reflectance data by compensating atmospheric absorption and scattering effects, we scanned three (white, silver, and black) 9 m × 9 m calibration tarps (Odyssey; Marlen Textiles, New Haven, MO, USA) using the PSR+ field spectrometer. We then incorporated the reflectance of the three calibration tarps with a radiative transfer model (MODTRAN 5.0; (Berk et al., 2008)) using a Bayesian method as previously documented (Wang et al., 2021).

To interpret the thermal image (details below) and understand metrics related to the ecosystem energy balance, we calculated vegetation indices including NDVI, VIS-NIR albedo, and the water band index (WBI; (Peñuelas et al., 1993)), using the airborne hyperspectral image.

Reflectance values at the 680 nm and 800 nm were used as the red and NIR bands to calculate NDVI. The water band index (WBI) was calculated as the ratio between reflectance at 900 nm and 970 nm. VIS-NIR albedo was calculated as mean reflectance in the 450 and 900 nm spectral range.

### Thermal data

The Jenoptik thermal camera (IR-TCM 640; Jena, Germany) had a broadband (7.5-14 μm) design. The field of view of the lens is 30°×23°, leading to a roughly 1.4 m ground pixel size. The measurement accuracy of the thermal camera was ±1.5°K within the range between 0°C and at-sensor radiance (*L_at-sensor_*). Georectification of thermal images were conducted using inflight 100°C. Lab based radiometric calibration was applied to the at-sensor DN value to calculate the position and posture information recorded by an independent GPS/IMU unit (NovAtel SPAN- CPT, Calgary, Alberta, Canada).

To derive the surface temperature from the airborne thermal image, we needed to solve two major issues: (1) compensating the atmospheric effects, and (2) separating emissivity and temperature. Theoretically, the relationship between at-sensor radiance, surface temperature and emissivity can be described as:

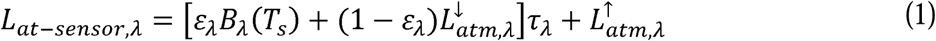

where *L_at-sensor_*_,λ_. is the radiance at wavelength *λ* measured at the sensor. *B*_λ_ (*T_s_*)i is the Planck radiation function at temperature T(K) and wavelength (λ). ε_λ_ is the emissivity of the target. τ*_λ_*is the atmospheric transmittance. 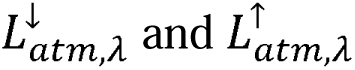 are the downward and upward atmosphere radiation.

We had one broad thermal band, thus we could only use the single-channel method for atmospheric correction and estimation of emissivity and surface temperature (Neinavaz et al., 2021, Li et al., 2013, Dash et al., 2002). Regarding single-channel method, a common way to apply atmospheric correction to thermal image is to rely on the radiative transfer models, such as MODTRAN. This strategy requires accurate estimation of air temperature, pressure and water vapor along the horizonal and vertical profile of the atmosphere, which was lacking in this study. Instead, we adapted an atmospheric correction method by including temperature measurements of the three calibration tarps using a handheld IRT (ThermoWorks, UT, USA) and thermocouple (Type T, Omega, CT, USA) linked to a datalogger (LI-100, LI-COR, NE, USA) on the ground.

According to the Stefan-Boltzmann law, the emissivity of the three calibration tarps were calculated as (T_IRT_/T_Thermocouple_)^4^. Mathematically, we can use three sets of ground measured emissivity and surface temperature of those calibration tarps to build three equations to directly solve the atmospheric variables 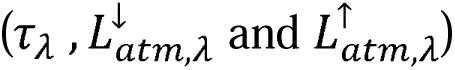 in equation (1). We then applied the atmospheric variables to the entire thermal image to obtain the ground level brightness temperature.

To calculate surface temperature from ground level brightness temperature, a simplified NDVI^THM^ method (Sobrino et al., 2008) was used to estimate per-pixel emissivity as:

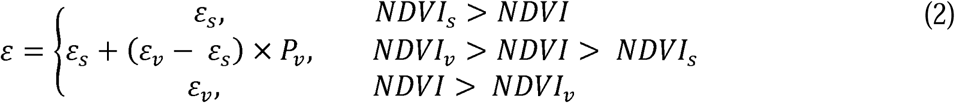

where *ε_s_* and *ε_v_* represented the emissivity of bare soil and dense vegetation, respectively. *Pv* denoted the percentage of vegetation coverage, which was calculated as:

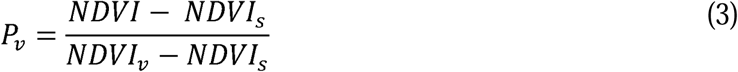

In this study, we used empirical emissivity values for dry sandy soil (*ε_s_*=0.95; (van Wijk, 1963)) This method maintained the continuity at NDVI = NDVIs (*Pv* = 0) and NDVI = NDVIv (*Pv* = 1). and vegetation (*ε_v_*=0.985; (Sobrino et al., 2008)). NDVI products were calculated using the hyperspectral image (details above). Because the spatial resolution of the hyperspectral images (1m) was finer than the thermal data (1.4m), for each pixel in the thermal image, the NDVI value of the pixel in the hyperspectral data with shortest distance was used.

## Results

Consistent effects of seeding and management treatments (burning and haying) emerged across multiple indicators of prairie productivity and energy balance from the grassland restoration experiment. Seeding and management treatments affected the percent cover of plants, litter, and bare soil, vegetation diversity (species and functional type composition) (Fig. 2). In general, the mid and high diversity plots had higher vegetation cover (Fig. 2a) and lower litter coverage (Fig. 2b) than the low diversity control plots. More species established successfully and consistently in the mid and high diversity plots, resulting in a lower percent cover of C_3_ grass (Fig. 2d), higher percent cover of C_4_ grass (Fig. 2e) and forbs (Fig. 2f), and higher overall plant diversity (Fig. 2g, h, i) than the control plots, in which two non-native species, including *Bromus inermis* and *Convolvulus arvensis* dominated. While there were no striking distinctions in species richness (Fig. 2g) or Shannon’s index (Fig. 2h&i) between the two seeding treatments, the high diversity plots had lower proportions of C_3_ plants (Fig. 2d) and higher proportions of C_4_ plants (Fig. 2e), leading to a higher functional diversity (Fig. 2i).

**Fig. 2.**
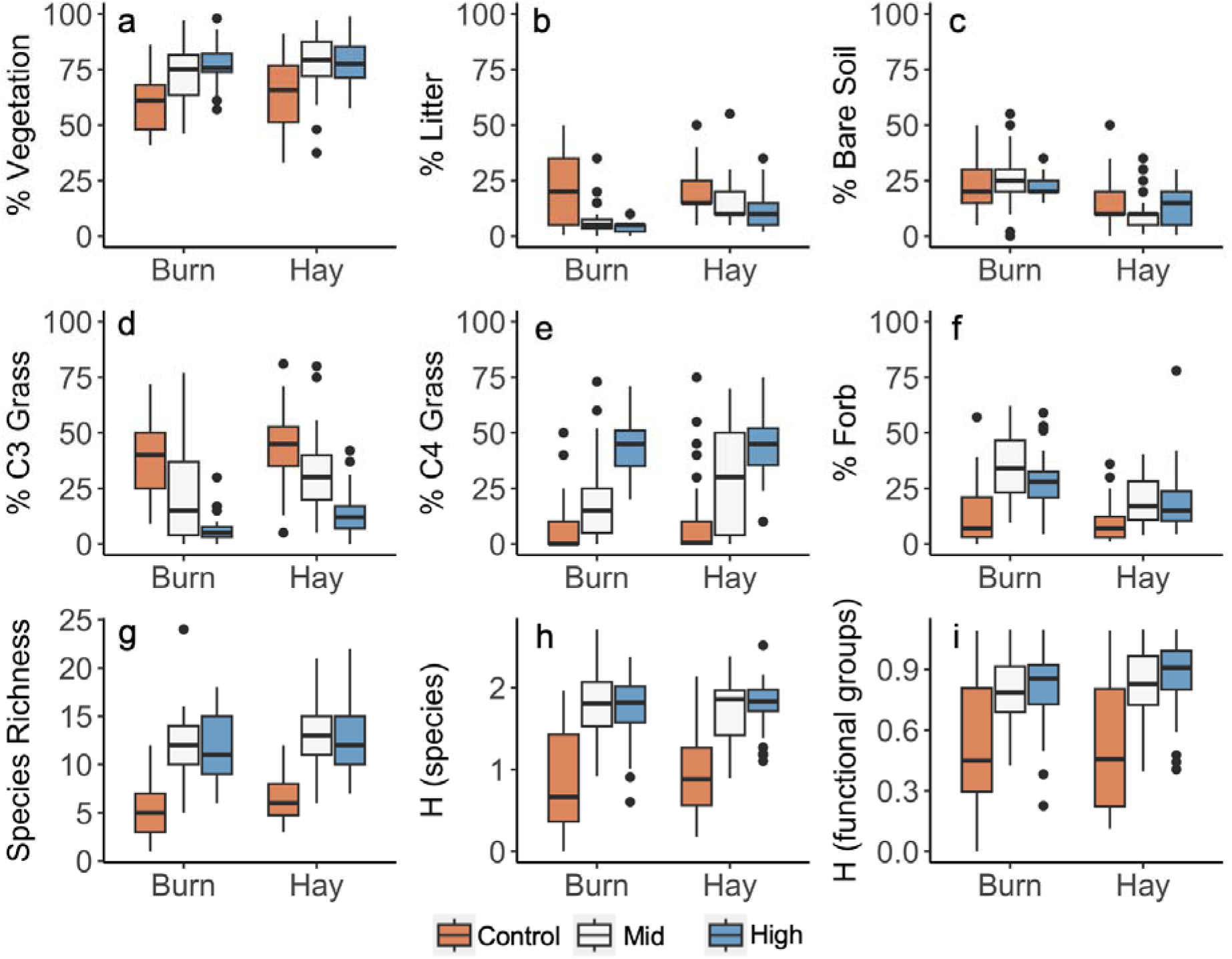
Effects of seeding treatments (control, medium, and high diversity) and managements (burn and hay) on percent coverage of vegetation (a), litter (b), bare soil (c), and three functional groups, including C_3_ grasses (d), C_4_ grasses (e), and forbs (f), species richness per m^2^ (g), Shannon’s index based on species (h), and Shannon’s index based on functional groups (i).

Burning removed dead plant material from previous growing seasons, resulting in reduced cover of litter in the mid and high diversity plots than hayed plots, but not in the control plots (Fig. 2b). The proportion of bare soil was also higher in the burned plots than hayed plots (Fig. 2c). Fire slightly suppressed the proportion of C_3_ grasses (Fig. 2d) and increased the proportion of forbs (Figure 2f), but led to no clear difference in C_4_ grass coverage (Fig. 2e). Within a burned block, the higher dominance of C_4_ within the high diversity seed mix was apparent. The mid diversity mix had some short and mid-stature native C_4_ and C_3_ grasses, but the tall, highly competitive C_4_ grasses (Indiangrass, big bluestem, and switchgrass) were excluded, which limited competition with forbs.

Ground and airborne spectroscopy clearly showed the effects of seeding and management (Fig. 3). Compared to the hayed plots, the burned plots exhibited a higher surface reflectance in the visible-near infrared spectral range (Fig. 2), although this was not clear in the ground sampling of the control plots (Fig. 3a), likely due to the small sample size. In the ground spectral measurements, which covered the full visible to shortwave infrared spectral range (400-2500 nm), the lignin and cellulose absorption features in the short-wave infrared were largely removed by burning (arrows; Fig. 3 b,c), indicating a reduction of standing dead biomass. This was more obvious in the mid (Fig. 3b) and high diversity (Fig. 3c) plots than the low diversity control plots (Fig. 3a) due to the higher vegetation coverage in these higher diversity plots. The airborne data captured the spectral difference (detected using ground measurements) among plots, but presumably with a more representative sample that measured the entire treatment plot area rather than the small subsample measured on the ground (Fig. 1). In particular, the airborne imagery recorded an increased albedo in burned plots (relative to hayed plots) and in the higher diversity plots (relative to the controls) (Fig. 3d,e,f).

**Fig. 3.**
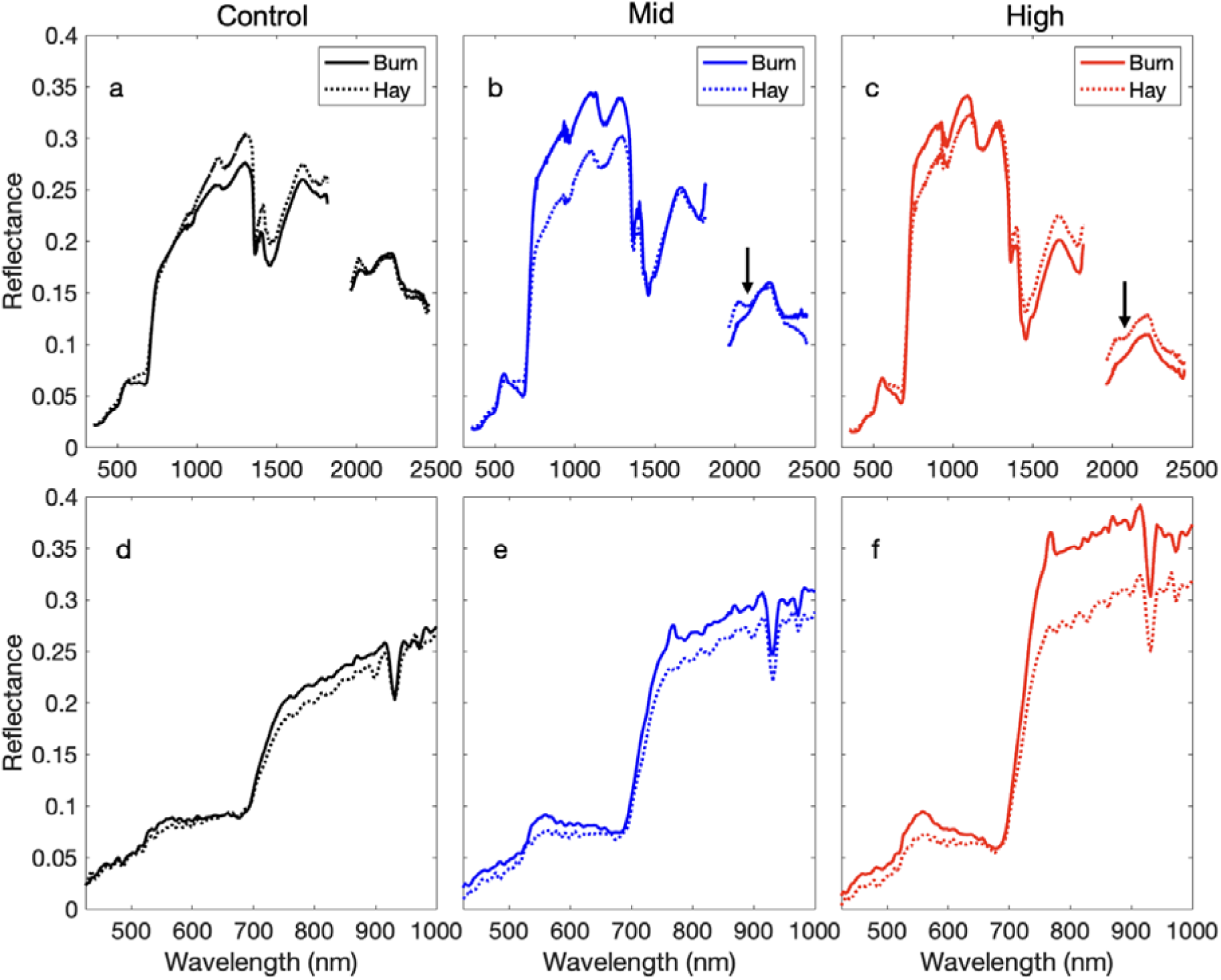
Mean reflectance spectra of transects from the six plots sampled from the ground (a, b, and c) and air (d, e, and f). Airborne data was extracted from the AISA Kestrel image using coordinates. The black arrow (panels b and c) indicates the lignin and cellulous absorption features that were missing from the burned plots. Note that the airborne data only covered the visible to near-infrared (400 – 1000 nm) spectral range. The ground and airborne spectra were plotted with different spectral ranges (X-axis scales).

The airborne imagery revealed complex spatial patterns of remote sensing indices related to productivity (NDVI), canopy water content (WBI), albedo (integrated reflectance) and surface temperature for the contrasting restoration treatments (Fig. 4). Higher NDVI, WBI, and albedo but lower temperature values (Fig. 4 a,b,c) generally corresponded to greener vegetation and more active plant growth, which in this case was also associated with greater plant species and functional diversity. These features were associated with large observed quantities of C_4_ grasses – mostly Indiangrass (*S. nutans*) with smaller amounts of big bluestem (*Andropogon gerardii* Vitman). Conversely, areas with low NDVI, WBI, and albedo (Fig. 4a,b,c), but high temperature (Fig. 4d) were areas dominated by C_3_ grasses, mostly *B. inermis* (smooth brome). The narrow, cool area with high NDVI roughly in the middle of the site (plot 13, Fig. 1a) was a low, wet area of extremely thick, tall C_4_ grass growth spanning the experiment, as plot 13 was a mid-diversity plot that was not disked and had heavy extant C_4_ grass cover prior to the experiment.

**Fig. 4.**
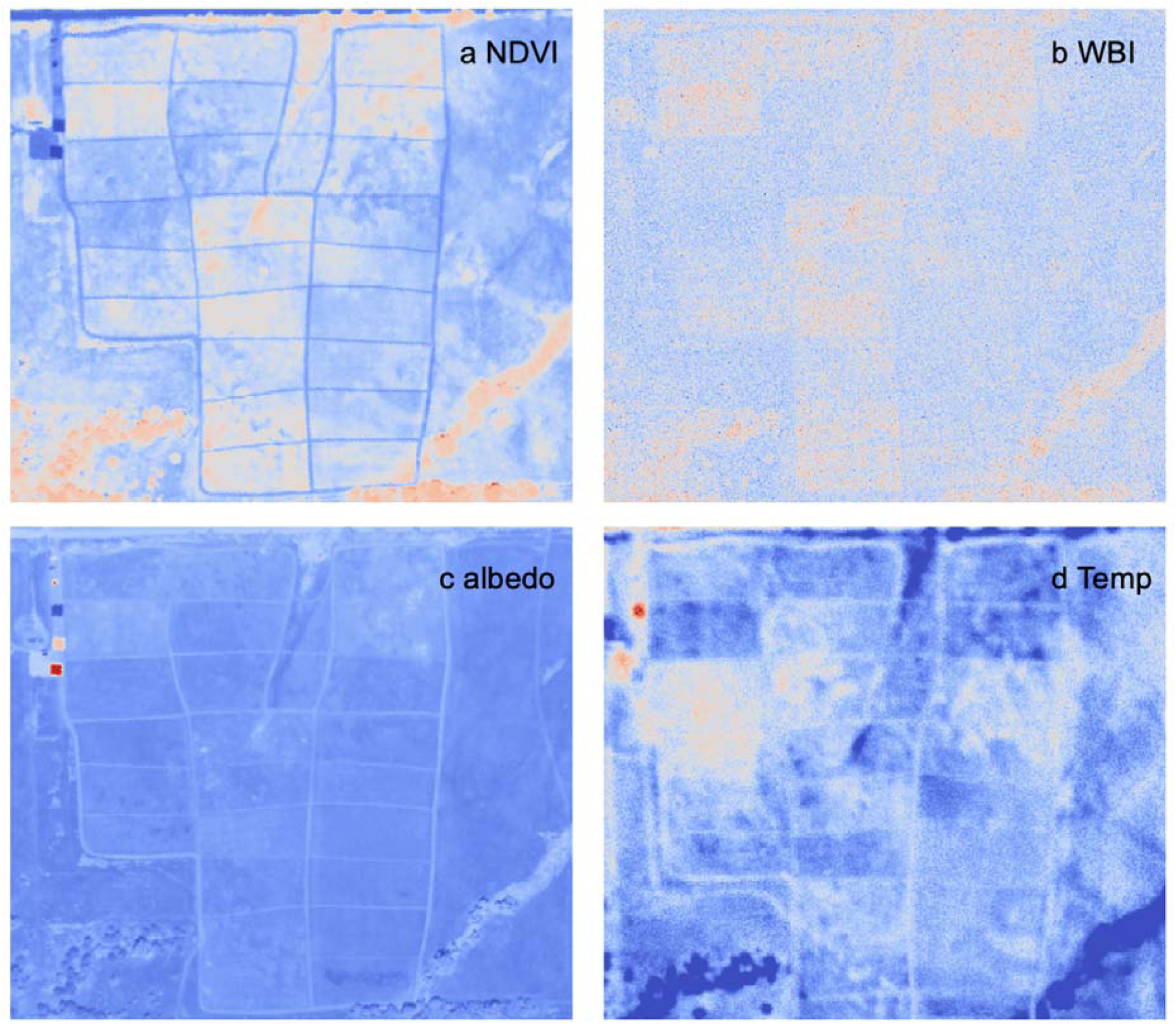
Airborne imagery revealing spatial patterns of indices related to surface energy balance, including productivity (NDVI; a), canopy water index (WBI; b), albedo (c), and surface temperature (d). For all panels, red indicates high values while blue indicates low values.

Overall, the airborne remote sensing products illustrated the combined effects of management (burning vs. haying) and seeding on productivity, vegetation water content and surface energy balance (Fig. 5). Burned plots had higher NDVI, WBI, and albedo and lower temperature than hayed plots and this pattern was more obvious in the mid and high diversity plots than the low diversity control plots (Fig. 5), indicating an interactive effect of seeding and management (P≤0.01).

**Fig. 5.**
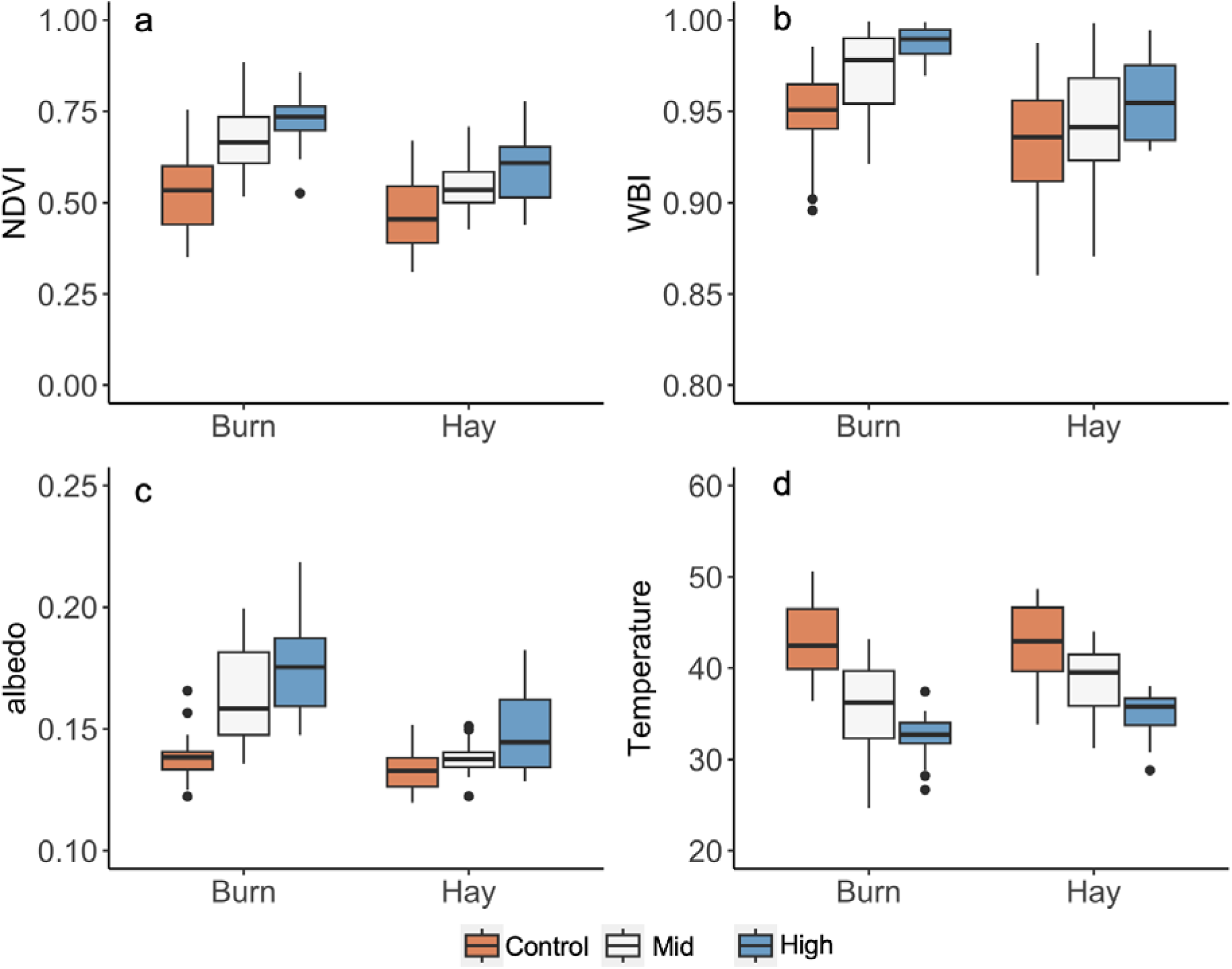
Responses of remote sensing indices related to surface energy balance, including productivity (NDVI; a), canopy water index (WBI; b), albedo (c), and Temperature (d) to seeding and management (burning and haying) treatments derived from airborne imagery.

Strong linear relationships were found between Shannon’s index based on species data (HSpecies) and vegetation indices and temperature calculated using airborne imagery (Fig. 6 and Table 3).

**Fig. 6.**
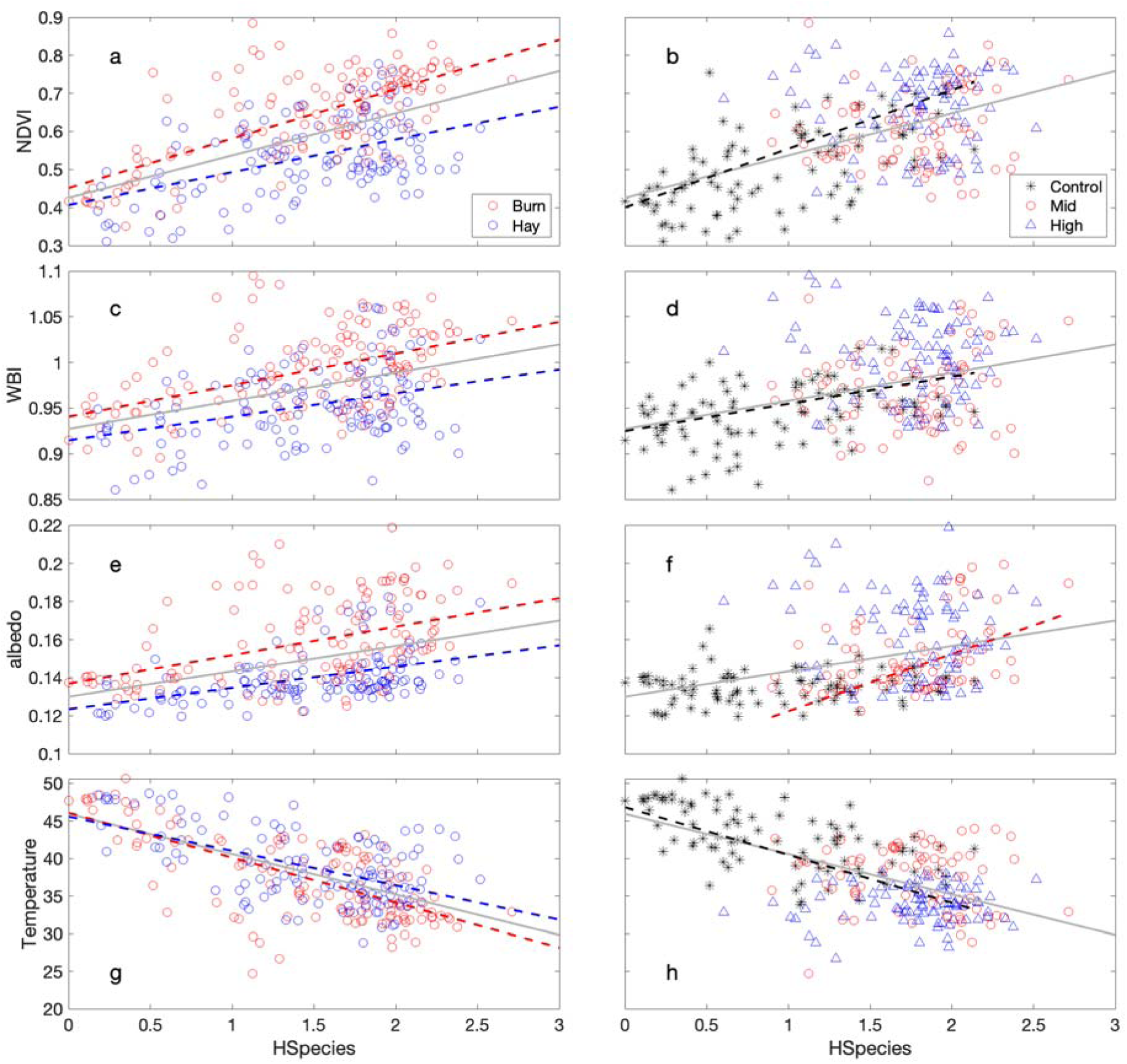
Relationships between Shannon’s index based on species (HSpecies) and various remote sensing products related to surface energy balance, including productivity (NDVI; a&b), canopy water index (WBI; c&d), albedo (e&f), and Temperature (g&h). Grey line indicates the relationship using all the data. Fit lines (dotted) are only shown for significant relationships (Table 3).

**Table 3.** Coefficient of determination (R^2^) of relationships between Shannon’s index based on species (HSpecies) and different remote sensing products. Data are combined (overall) and split by management (burn vs hay) and seeding treatments (control, mid and high diversity). Significant codes: NS, 0.05 < p, *, 0.01 < p < 0.05, **, 0.001 < p < 0.01 and ***, p < 0.001.

| Indices | Overall | Management |  | Seeding |  |  |
| --- | --- | --- | --- | --- | --- | --- |
|  |  | Burn | Hay | Control | Mid | High |
| NDVI | 0.297*** | 0.483*** | 0.239*** | 0.336*** | NS | NS |
| WBI | 0.158*** | 0.246*** | 0.13*** | 0.135*** | NS | NS |
| albedo | 0.153*** | 0.19*** | 0.224*** | NS | 0.127** | NS |
| Temperature | 0.39*** | 0.471*** | 0.309*** | 0.382*** | NS | NS |

These relationships varied with management (Fig. 6 a,c,e,g) and seeding (Fig. 6 b,d,f,h) treatments. Burning affected the slopes of the NDVI-Shannon’s index (P=0.01) and the intercept of the WBI-Shannon’s index and the albedo-Shannon’s index (P<0.05). Neither slope, nor intercept of the temperature-Shannon’s index relationship was significantly different between the burned and hayed plots. When data were split by seeding treatments, there was no significant relationship between remotely sensed data and Shannon’s index for the mid and high diversity plots, except for the relationship between albedo and Shannon’s index for the mid diversity plots (Fig. 6 f). For the control plots, significant relationships were found between NDVI, WBI, temperature and Shannon’s index (Fig. 6 b,d,h).

Comparing the remote sensing products to the percent coverage of C_3_ grasses further revealed clear effects of functional type composition on remotely sensed data (Fig. 7 and Table 4).

**Fig. 7.**
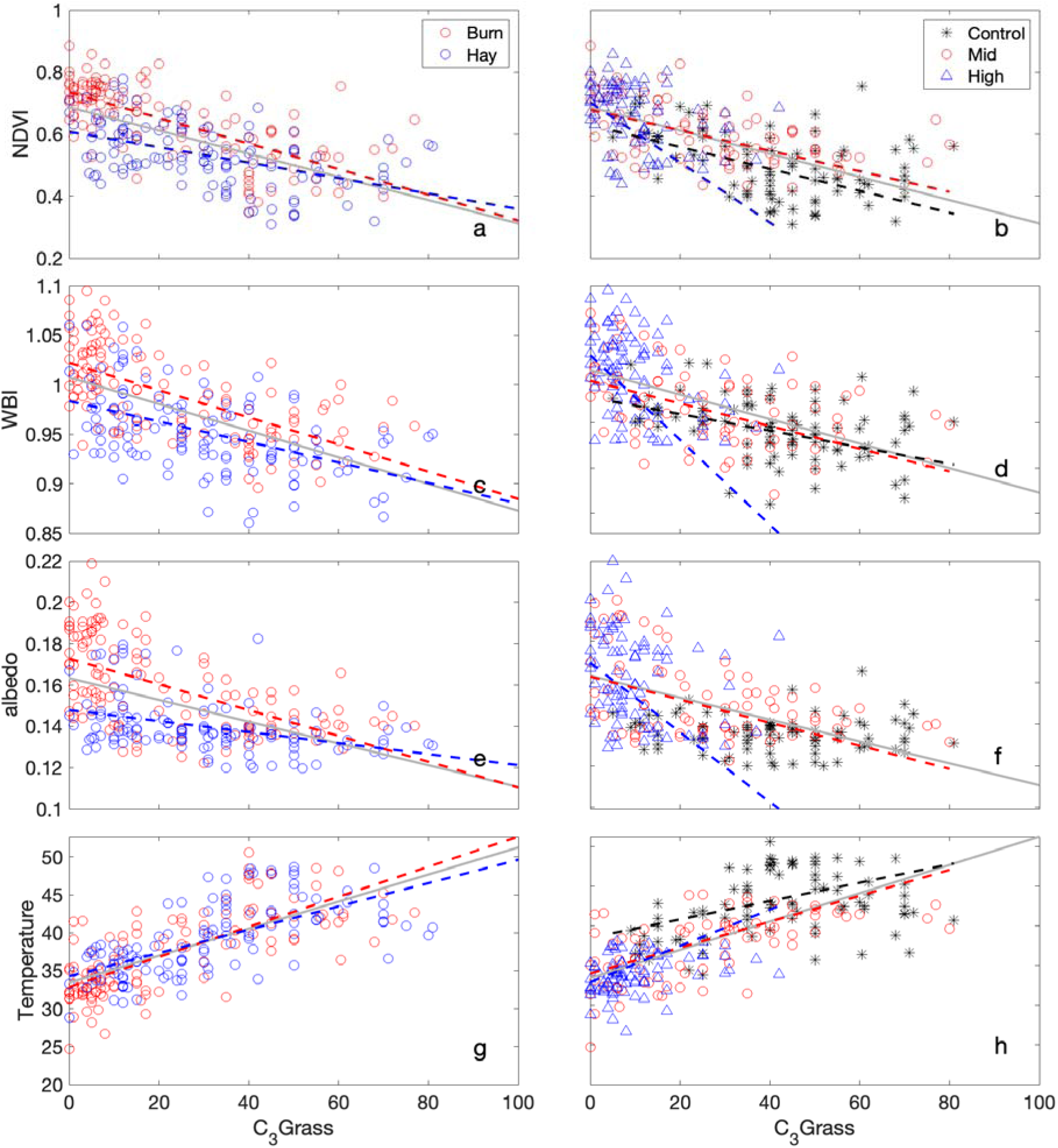
Relationships between percent coverage of C_3_ grass and various remote sensing products related to surface energy balance, including productivity (NDVI; a&b), canopy water index (WBI; c&d), albedo (e&f), and Temperature (g&h). Grey line indicates the relationship using all the data. Fit lines (dotted) are only shown for significant relationships (Table 4).

**Table 4.** Coefficient of determination (R^2^) of relationships between percentage cover of C_3_ grasses and different remote sensing products. Data are combined (overall) and split by management (burn vs hay) and seeding treatments (control, mid and high diversity). Relationship with *P* value > 0.05 is shown as NS.

| Indices | Overall | Management | Seeding |
| --- | --- | --- | --- |

|  |  | Burn | Hay | Control | Mid | High |
| --- | --- | --- | --- | --- | --- | --- |
| NDVI | 0.394 | 0.516 | 0.245 | 0.198 | 0.27 | 0.128 |
| WBI | 0.352 | 0.41 | 0.254 | 0.142 | 0.207 | 0.162 |
| albedo | 0.279 | 0.35 | 0.155 | NS | 0.208 | 0.08 |
| Temperature | 0.5 | 0.546 | 0.422 | 0.142 | 0.37 | 0.127 |

Significant differences in the intercepts, but not slopes, of productivity (NDVI)-C_3_ percent cover and canopy water content (WBI)-C_3_ percent cover were found between the burned and hayed plots. For the albedo-C_3_ percent cover relationship, both slopes and intercepts were significantly different (P<0.001) between the burned and hayed plots. There was no significant difference in slopes or intercepts between the burned and hayed plots in the temperature-C_3_ percent cover relationship. For the seeding effects, the high diversity plots were different from the mid and low diversity plots in the productivity-, canopy water content-, and albedo-C_3_ percent cover relationships, which was presumably caused by the relatively low percent cover of C_3_ grasses in the high diversity plots. However, no difference was noticed in the temperature-C_3_ percent cover relationships among the seeding treatments.

## Discussion

### Links between diversity, productivity and energy balance

In grasslands, the relationship between biodiversity, ecosystem function, and energy balance is complex and multifaceted, but key findings have emerged, suggesting co-benefits of certain management choices. In general, higher diversity grasslands tend to have higher productivity, expressed as higher vegetation percent coverage and density and more complex leaf traits and 3D canopy structure, which can together lower the surface temperature by affecting both energy absorption, reflection, evapotranspiration and sensible heat flux (Guimarães-Steinicke et al., 2021, Milcu et al., 2016, Vojtech et al., 2008). In this study, the high vegetation coverage in the high diversity plots likely had higher thermal mass and evapotranspiration (suggested by the higher canopy water content), lowering the surface temperature (Figs 4 and 5). Additionally, the higher albedo of the higher diversity plots likely contributed to a greater cooling effect and lower surface temperature.

The composition of species or functional groups rather than species richness *per se* affects the energy balance in grasslands (Petersen and Isselstein, 2015, Schouten, 2022). Under high soil moisture conditions, C_3_ grass areas would be cooler than C_4_ grass areas because C_3_ grasses have higher stomatal conductance and transpiration for the same amount of leaf productivity, lowering surface temperature. However, this situation likely changes with seasonal conditions: the low temperature values in our high diversity plots directly corresponded to the high dominance of tall C_4_ grasses, especially when burned (Fig. 2). This is probably because our data was collected in the mid growing season when the cool season grasses were seasonally inactive before greening up again in the fall. Also, due to the relatively dry and hot summer in our field site, little soil moisture was available for the shallow-rooted smooth brome grass and their stomatal conductance and transpiration had largely shut down at the point when we took measurements. In contrast, the deeper rooted native C_4_ grasses like big bluestem remained green and were able to maintain their higher stomatal conductance and transpiration led to evaporative cooling of the vegetation surface. These observations suggest that the direction and magnitude of the temperature effect may differ under different conditions or time of year.

### Fire/hay treatment modulates these effects

Management regimes (burning and haying) affect grassland diversity and modulate the coupling between vegetation and the atmosphere in complex ways (Gordijn and O’Connor, 2021).

Phylogenetic lineage and lifecycle rather than photosynthetic type may determine the response of different species to fire, and this response is further affected by the frequency and timing of fire (Ripley et al. 2015). In this study, the prescribed burn occurred when the smooth brome (*Bromus inermis*) was fully grown but had not yet set seed, and thus may have temporarily exhausted the plant’s resources for further growth that year. Also, increased soil temperature and evapotranspiration associated with the post-burn conditions would also have aggravated smooth brome in a drought year and encouraged it to shut down transpiration mid-growing season. By removing dead thatch from previous seasons, suppressing C_3_ grass (mostly the smooth brome) growth, the prescribed burn stimulated native C_4_ grass growth, possibly because shade intolerant C_4_ grasses can grow faster than C_3_ grasses in hot and open conditions and benefit from biomass removal (Osborne and Beerling, 2005). These observations indicate that timing of particular management treatments may be critical in attaining key management outcomes.

### The role of grassland restoration program in climate mitigation

Ecological restoration is an effective solution to counter the loss of biodiversity and ecosystem services, and also offers a powerful tool to mitigate climate change. Through energy balance (albedo and evapotranspiration), carbon uptake and storage, and other processes, terrestrial ecosystems, including grasslands, can amplify or dampen climate change rising from anthropogenic greenhouse gas emission. Similar effects of productivity and energy balance have been noted for forests (e.g., Bonan 2008), but have generally not been considered for grasslands, which cover roughly 40 percent of terrestrial surfaces and dominate the interannual variability in the terrestrial carbon sink (Wang et al., 2022, Ahlström et al., 2015, Poulter et al., 2014), offering significant climate mitigation potential. In our study, these effects were detectable in the altered productivity, albedo and surface temperature in treatment plots (Figs 2, 4, and 5). While we detected enhanced productivity in more diverse plots, we did not test below-ground carbon directly but numerous studies from experimental grasslands suggest that enhanced diversity and above-ground productivity can, over time, enhance soil carbon storage (Lange et al., 2015, Steinbeiss et al., 2008). We expect that restoring diverse productive grasslands can reduce atmospheric CO_2_ content by storing carbon biomass, particularly below-ground biomass.

However, while restoration activities may start the recovery process relatively quickly, a full recovery of a former community and the below ground structure of a secondary grassland may take longer (Nerlekar and Veldman, 2020, Weisser et al., 2017). For this reason, we recommend long-term studies of grassland restoration practices, not only considering diversity and productivity, but also surface-atmosphere feedbacks in the context of various management treatments that include grazing and burning, which have been an integral part of historical prairie management.

## Acknowledgements

Funding: NSF and NASA grant (DEB-1342823); University of Nebraska Collaboration Initiative Grant; Nebraska Environmental Trust (Prairie Corridor Phase III, #16-122)

## Author contributions

Conceptualization: R.W., J.A.G., D.A.W.; Methodology: R.W., J.A.G., K.F.E.H., P.R.K., D.A.W.; Investigation: R.W., J.A.G., K.F.E.H., P.R.K., D.A.W.; Writing–original draft: R.W., J.A.G., K.F.E.H.; Writing–review and editing: R.W., J.A.G., K.F.E.H., P.R.K., D.A.W.

## Data availability

data used in this work will be made available through the University of Nebraska research data repository (SANDY).

